# High-throughput Phenotyping with Temporal Sequences

**DOI:** 10.1101/590307

**Authors:** Hossein Estiri, Zachary H Strasser, Shawn N. Murphy

## Abstract

**Objective:** High-throughput electronic phenotyping algorithms can accelerate translational research using data from electronic health record (EHR) systems. The temporal information buried in EHRs are often underutilized in developing computational phenotypic definitions. The objective of this study is to develop a high-throughput phenotyping method, leveraging temporal sequential patterns of discrete events from electronic health records.

**Materials and Methods:** We develop a representation mining algorithm to extract five classes of representations from EHR diagnosis and medication records: the aggregated vector of the records (AVR), the traditional immediate sequential patterns (SPM), the transitive sequential patterns (tSPM), as well as two hybrid classes of SPM+AVR and tSPM+AVR. A final small set of representations were selected from each class using the MSMR dimensionality reduction algorithm. Using EHR data on 10 phenotypes from Mass General Brigham Biobank, we trained regularized logistic regression algorithms, which we validated using labeled data.

**Results:** Phenotyping with temporal sequences resulted in a superior classification performance across all 10 phenotypes compared with the AVR representations that are conventionally used in electronic phenotyping. Although this study only utilizes the diagnosis and medication records, the high-throughput algorithm’s classification performance was superior or similar to the performance of previously published electronic phenotyping algorithms. We characterize and evaluate the top transitive sequences of diagnosis records paired with the records of risk factors, symptoms, complications, medications, or vaccinations.

**Discussion:** The proposed high-throughput phenotyping approach enables seamless discovery of sequential record combinations that may be difficult to assume from raw EHR data. A transitive sequence can offer a more accurate characterization of the phenotype, compared with its individual components. Additionally, the identified transitive sequences of a given phenotype reflect the actual lived experiences of the patients with that particular disease.

**Conclusion:** Sequential data representations provide a precise mechanism for incorporating raw EHR records into downstream Machine Learning.

## INTRODUCTION

Biomedical researchers are progressively applying modern Machine Learning (ML) algorithms to data from electronic health records (EHRs). Despite the natural excitement about the large amount of information presented by EHRs, daunting challenges remain. As the primary impetus for EHR implementation has been clinical care, EHR observations reflect a complex set of processes that further obscure their utility in research. Dimensionality, sparsity, heterogeneity, and quality issues present significant impediments for secondary use of EHR data.[1,2] In particular, the EHR observation records are often not direct indicators of a patient’s true health state, but rather reflect the clinical processes (e.g., policies and workflows of the provider and payor organizations), the patient’s interaction with the system, and the recording process.[3–5]

As a result of the biases inherent in EHR data, identifying cohorts of patients with certain health conditions can become complex. In order to make precise assumptions about the presence of a disease, we would need to perform phenotyping. The key task in phenotyping is to identify patient cohorts with (or without) a certain phenotype or clinical condition of interest.[6,7] Developing specialized phenotypic definitions from EHR data can be expensive and often requires involvement by domain experts. [8–11] Nevertheless, due to its critical role in re-using EHR data from research, several healthcare institutions are actively involved in constructing and validating electronic phenotyping algorithms. Efforts to curate computational phenotypes and discover clinical knowledge from EHR observations must account for the potential biases introduced through the recording process.[4]

Another underutilized aspect of electronic medical records is their temporal dimension. EHRs contain important temporal information about disease progression and treatment outcomes. However, EHR observations are often acquired asynchronously across time (i.e., measured at different time instants and sampled irregularly in time) and include sparse and heterogeneous data.[12–17] These properties challenge the application of standard temporal analysis methods to clinical data recorded in EHRs.

The record of the EHR diagnosis and its time stamp may not give the true disease state or the actual onset of the disease. In this paper, we utilize a novel sequential pattern mining algorithm to construct temporal data representations from EHR data. Using a high-throughput feature selection algorithm, we then utilize the temporal sequential representations to develop computational phenotyping algorithms for 10 phenotypes. We demonstrate that the temporal sequential features significantly outperform raw EHR features that are commonly used in computational phenotyping algorithms.

## BACKGROUND

Although electronic health records often include incomplete, inaccurate, or even biased data, the wealth of information they provide is sufficient for constructing clinically-relevant sets of observable characteristics that define a disease or phenotype.[4,18] The task in electronic phenotyping is to determine patients with (or without) certain phenotypic characteristics based on data from electronic medical records,[7] which is challenging due to the heterogeneity and complexity of multimodal EHR data.[6] As a result, developing specialized phenotypic definitions from EHR data is generally expensive.[8,9]

Approaches to electronic phenotyping include rule-based methods and computational methods, which can be broadly characterized under text processing and supervised/semi-supervised/unsupervised statistical learning techniques.[6,19] Applying supervised and semi-supervised machine learning to EHR data for identifying cohorts with clinical phenotypes is rapidly prevailing.[11,19–37] Although, such phenotyping models require a human-annotated gold-standard training set, which remains a bottleneck.[30] In addition, defining clinically-meaningful EHR features for computational phenotyping relies on a heavy dose of domain expert involvement, using complex ad-hoc procedures that are often hard to generalize and scale.[8–11] Despite the cost, several academic medical centers are actively involved in constructing and validating EHR phenotyping algorithms. Some of the notable efforts include the i2b2-centered efforts led by Harvard University and Mass General Brigham,[20,21,31–37] the BioVU led by Vanderbilt University,[22,23] and the multicenter eMERGE Network consortium,[24–27] the Phenotype Knowledgebase website, PheKB,[28] and the OMOP-centered Automated PHenotype Routine for Observational Definition, Identification, Training and Evaluation (APHRODITE)[38], to name a few.

For developing computational phenotyping algorithms with EHR data, features are typically constructed by identifying relevant clinical events (e.g., diagnoses or medication records from structured data or certain keywords from the clinical notes). A vector of these records is obtained from patient-level marginal counts and cohort-level aggregation. This approach is rudimentary and misses potentially useful information that is available in the electronic medical records. Of note, EHRs provide a wealth of longitudinal information that can be leveraged to improve computational phenotyping algorithms.[18] Temporal representation mining methods offer technical solutions that can account for this aspect of the EHR.

Temporal representation mining involves providing a machine-readable representation that formalizes the concept of time as it is relates to a set of events and temporal relationships.[39,40] In biomedical research, development and evolution of temporal representation mining approaches has been largely focused on the temporal abstraction (TA)[41] of data from continuous clinical measurements.[42] Yet, discrete clinical data, such as diagnoses, medications, and procedures, are poor candidates for the TA approach. While numerical observations, such as laboratory test results, have explicit timestamps, precise numeric time-stamped information is often unavailable in discrete clinical data.

Sequential Pattern Mining (SPM)[43] is a viable alternative approach for discrete data. The goal in SPM is to discover “relevant” sub-sequences from a large set of sequences (of events or items) with time constraints. The relevance is often determined by a user-specified occurrence frequency, known as the minimum support.[44] The frequent sequential pattern (FSP) problem is to find the frequent sequences among all sequences.[45] A priori-based sequential pattern mining methods, such as sequential pattern discovery using equivalence classes (SPADE)[46] and sequential pattern mining (SPAM),[47] are popular in the healthcare domain. For example, Perer,Wang, and Hu (2015) used the SPAM algorithm for mining long sequences of events. [25] The a priori property is that if a sequence cannot pass the minimum support test (i.e., is not assumed frequent), all of its sub-sequences will be ignored. However, for various reasons, temporal patterns that are mined based on frequency may not make clinical sense. For example, low blood pressure readings after the administration of a specific, but irrelevant, medication may be frequently observed, yet have no real clinical meaning.[48]

Recently, we introduced the transitive sequential pattern mining (tSPM) algorithm along with an early implementation of the MSMR algorithm.[49] The tSPM algorithm provides a modified sequencing procedure to address some of the issues caused by the recording processes in EHRs. The MSMR algorithm, offers a high-throughput feature engineering technique to improve the frequency-based a priori property in the traditional SPM approach. As a proof-of-concept, we showed that the sequencing approach improves disease prediction and classification in a single disease. In this study, we apply the tSPM algorithm – with an improved MSMR algorithm, – to computational phenotyping in EHR data. We perform a comprehensive comparison of the transitive and traditional sequential representations with the conventional way of using EHR observations as features for computational phenotyping in 10 phenotypes. We also compare the phenotyping performance with the state-of-the-art phenotyping algorithms published in research informatics literature.

## MATERIALS AND METHODS

This study aims to answer a principle question: can temporal representations mined through sequential pattern mining improve computational phenotyping with EHR data? Developing the computational phenotyping algorithms with temporal sequences follows two steps: 1) a phenotype-agnostic representation mining and 2) a semi-supervised phenotype-specific dimensionality reduction, the MSMR algorithm (Figure 1). We use terms feature and data representation interchangeably.

**Figure 1.**
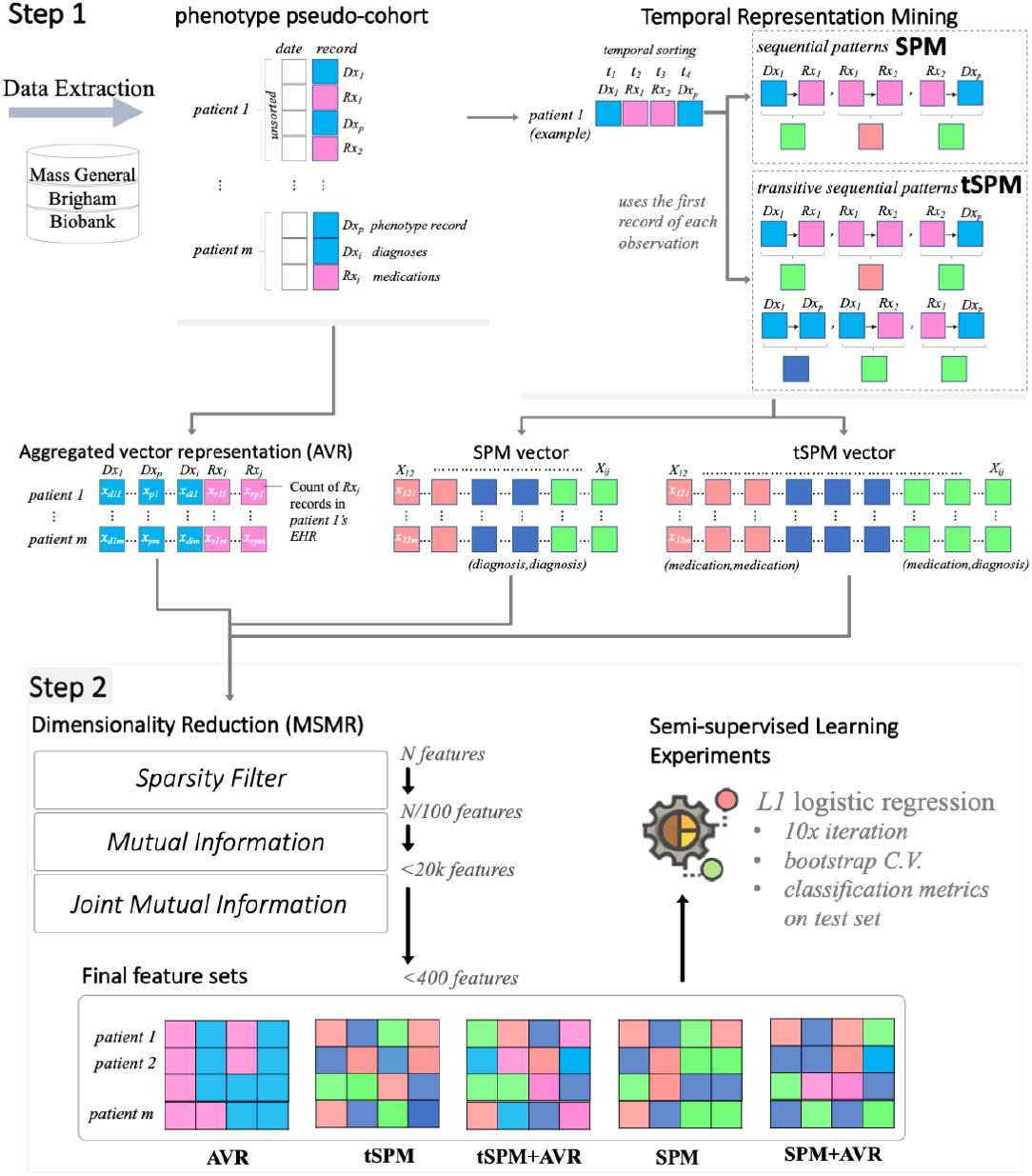
The two steps involved in the high-throughput phenotyping with temporal sequences.

### Representation mining

We only used the medication and diagnosis records data. For the diagnosis records, we used the International Classification of Diseases, ninth/tenth Revision, Clinical Modification (ICD 9/10 CM). For medications, we use RxNorm codes. Given a list {*R*_1_, *R*_2_, *…, R*_n_} of diagnosis or medication records, for each patient *p*, we recorded the times 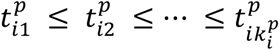 at which the record *R*_*i*_ was logged (we allow 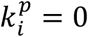, in which case record *R*_*i*_ was yet to be recorded for patient *p*). We then mined three vectors of data representations:

First, we constructed a baseline representation that applies the conventional approach for using electronic health records as features for computational phenotyping. We will henceforth call this the Aggregated Vector Representation (AVR) approach.

Second, we mined a set of temporal sequential representations by sequencing the medication and diagnosis records in electronic medical records. For the temporal sequencing, we utilized the traditional Sequential Pattern Mining schema (SPM), in which immediate sequences are mined.

Third, and to account for irregularity of clinical records and the recording processes, we mined the novel transitive Sequential Pattern Mining schema (tSPM),[49] in which sequences of unlimited lengths are possible.

#### AVR representations

In the AVR approach, which is the conventional approach for using EHR data as records for computational phenotyping, the marginal count of a vector of a selected medical record (often diagnosis codes) is calculated for each patient. The patient is represented by a vector of the length equal to the number of unique events in her/his medical records. The initial set of AVR representations are all possible records, and for each patient, we record only the numbers 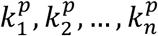 of each record. For each *i*, we think of the 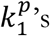 as samples of a random variable *X*_*i*_. Our goal is then to predict the class label *Y*, given *X*_1_, *X*_2_, …, *X*_3_.

For example, when Type 2 Diabetes Mellitus (T2DM) is a feature in the AVR approach, the number of times the diagnosis record for T2DM is recorded in a patient’s electronic record is used as the classifier for training and testing.

#### SPM representations

In the traditional SPM approach, the features are all possible *pairs* of distinct records (*R*_*i*_, *R*_*j*_), *i* ≠ *j*. For a given patient *p* and a given time *t*, let *t*^*’*^ > *t* be minimal such that for some *ℓ*≤ and some 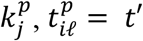 That is, *t′* is the first time strictly bigger than *t* at which some observation is made for the patient. For a given patient *p*, and a given index *i ∈* {1,2, …, *n*}, let *S*_*pi*_ be the set of all pairs (*j*, ℓ′), with *j ∈* {1,2, …, *n*} and 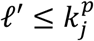, such that the record *j* is logged immediately after record *i* at time 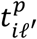 Formally, (*j, ℓ′*) *∈ S*_*pi*_ if and only if 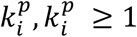 and there exists 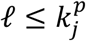 such that (*t*_*iℓ*_)′ = *t*_*iℓ*_′

For each *i, j ≤ n, i* ≠ *j*, let *r*_*ijp*_ = |*S*_*ip*_*|*. We think of the *r*_*ijp*_’s as samples of a random variable *X*_*ij*_ and the goal is to predict the class label *Y* given 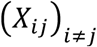.

#### tSPM representations

In the transitive sequential pattern mining (tSPM) algorithm, the features are again all possible *pairs* of distinct medical records (*R*_*i*_, *R*_*j*_) *i ≠ j*. For a fixed patient *p*, and *i ≠ j ≤ n*, we set *r*_*ijp*_ to be 1 if 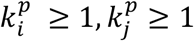, *and* 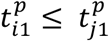0 otherwise. In words, *r*_*ijp*_ is 1 if and only if both records *R*_*i*_ and *R*_*j*_ were logged for the patient, and the *first* record of medical record *i* was before, or at the same time as, the first record of medical record *j*. Here, for each fixed *i ≠ j*, we think of the *r*_*ijp*_’s as samples of a random variable *tX*_*ij*_. Then our goal is to predict the class label *Y* given 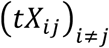. The use of the first record (rather than all records) is a specification difference in the way the sequential patterns are organized in tSPM compared to SPM. This difference helps to address the issue of repeated problem list entries.[49]

### Dimensionality reduction

If all pairs of sequences in the transitive sequencing approach exist, there will be exactly 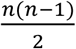 pairs (*i, j*) with *i ≠ j* and *i, j ≤ n*. To extract features for phenotyping, we applied a formal dimensionality reduction procedure that aims to minimize sparsity and maximize relevance (MSMR)[49] to all three feature vectors. To minimize sparsity, MSMR removes any feature that has a prevalence smaller than 0.5 percent. For maximizing relevance, MSMR is principally a semi-supervised dimensionality reduction algorithm that takes a silver-standard class label *Y’* to compute information gain metrics for all features. MSMR is able to effectively scale to large dimensionality spaces, and thus is a high-throughput algorithm.

For the remaining features, MSMR computes the empirical mutual information using an estimation of the entropy of the empirical probability distribution.[50,51] Mutual information provides a measurement of the mutual dependence between two random variables, which unlike most correlation measures can capture non-linear relationships.[51,52] We ranked the data representations based on the computed mutual information with the silver-standard labeled outcome (in ties, we used prevalence to determine the ranking) and conventionally select the top 20,000 data representations from each approaches.

We further dissected the relevance property by applying a filter-type feature extraction method using joint mutual information (JMI).[53] The algorithm starts with a set *S* containing the top feature according to mutual information, then iteratively adds to *S* the feature *X* maximizing the *joint mutual information score*:

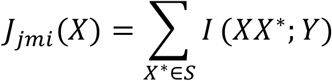

Here, *I*(*Z*; *Y*) denotes the mutual information between random variables *Z* and *Y* (a measure of the information shared by *Z* and *Y* — it can be expressed as the entropy of *Z* minus the entropy of *Z* given *Y*). The random variable *XX*^*\**^ is simply the random variable corresponding to the joint distribution of *X* and *X*^*\**^. In the end, we select the top features that were added to the set *S*. The joint mutual information score also takes into account the redundancy between the features: two features could each be highly relevant on their own, but also be strongly correlated. Brown et al (2012) suggested that the JMI score provides the “best trade-off […] of accuracy and stability.”[54]

The JMI step allows for integration of representations from different types. To obtain combined feature sets (AVR + sequential patterns), we also compute the joint mutual information when the AVR and SPM/tSPM representations are available.

### Study populations

We used data on 10 phenotypes from the Mass General Brigham (MGB) Biobank: Alzheimer’s disease (AD), chronic obstructive pulmonary disease (COPD), congestive heart failure (CHF), coronary artery disease (CAD), stroke, rheumatoid arthritis (RA), type I diabetes mellitus (T1DM), type II diabetes mellitus (T2DM), ulcerative colitis (UC), and atrial fibrillation (AFIB). For each phenotype, two pseudo-cohorts are available. Based on a list of ICD-9/10 codes for the respective phenotype, a given patient in the pseudo-cohort has at least one record of the respective diagnosis code(s), as well as an outcome label, which determines the “true” presence of the phenotype. We use the term “pseudo-cohort” to distinguish these patient cohorts from validated disease cohorts.

For each of the phenotypes, a small dataset included patient pseudo-cohorts with gold-standard outcome labels. To create gold-standard labels, teams of board-certified clinicians and/or nurses reviewed clinical notes and other required data of samples of patients selected from patients consented into the MGB Biobank between April 2012 and April 2017 and with 1 or more diagnosis record of the phenotype. The gold-standard phenotype pseudo-cohort datasets include labels for an average of 351 patients (ranging from 163 to 700 patients). For the 10 phenotypes, larger pseudo-cohorts were also pulled from the Biobank, with an average population of over 6,000 patients. For these patients, silver-standard labels are curated using a generative transfer learning algorithm. [cite: 10.1093/jamia/ocaa215] The average number of phenotype records was 28 in the gold-standard datasets and 20 in the silver-standard datasets. Table 1S (in supplementary materials) provides descriptive data on each of the phenotype pseudo-cohorts. The use of data for this study was approved by the Mass General Brigham Institutional Review Board (2017P000282).

**Table 1.** The number of unique representations by type before and after MSMR’s sparsity reduction.

| phenotype | mined unique representations |  |  | representations after sparsity screening |  |  |
| --- | --- | --- | --- | --- | --- | --- |
|  | *AVR* | *SPM* | *tSPM* | *AVR* | *SPM* | *tSPM* |
| **AD** | 6,193 | 209,389 | 3,844,039 | 1,661 | 28,672 | 1,211,853 |
| **AFIB** | 9,050 | 172,134 | 3,190,557 | 5,460 | 23,957 | 795,316 |
| **CAD** | 14,406 | 476,617 | 9,142,990 | 5,599 | 23,480 | 810,862 |
| **CHF** | 6,857 | 391,679 | 7,419,996 | 6,415 | 39,265 | 1,349,704 |
| **COPD** | 6,284 | 369,062 | 6,527,854 | 5,412 | 37,721 | 1,556,750 |
| **RA** | 5,203 | 305,787 | 5,782,618 | 4,517 | 19,157 | 790,214 |
| **Stroke** | 5,573 | 281,298 | 4,692,549 | 3,395 | 24,186 | 839,268 |
| **T1DM** | 6,673 | 385,626 | 7,086,684 | 3,334 | 46,260 | 1,538,308 |
| **T2DM** | 10,439 | 429,115 | 8,376,105 | 5,887 | 26,331 | 920,251 |
| **UC** | 4,904 | 207,256 | 3,920,849 | 1,893 | 14,651 | 603,902 |

### Model Training and Evaluation

We applied logistic regression classifiers with *L1* regularization to the training sets (with silver-standard labels) for developing the computational phenotyping algorithms, using bootstrap cross-validation. Regularized logistic regression classifiers are the most popular classifiers in EHR phenotyping – for example, see [8,11,38,55–57]. From each data representation class (AVR, SPM, and tSPM), a nested set of the top 50 to 200 representations are extracted from the MSMR algorithm for phenotyping. In addition, using the MSMR algorithm, we extracted hybrid feature sets that included both the AVR and sequential representations. This resulted in two additional feature sets combining the top AVR representations with the SPM and tSPM representations. Overall, we trained computational phenotyping algorithms on five classes of feature sets: 1) AVR, 2) SPM, 3) tSPM, 4) AVR+SPM, and 5) AVR+tSPM.

We evaluated the phenotyping algorithms against the gold-standard labels available in the held-out test sets to compute the area under the Receiver Operating Characteristic (ROC) curve (AUC ROC). Furthermore, we iterated the training process 10 times with bootstrap sampling and use the median performance metrics for comparing the feature sets. Overall, for each phenotype, we trained 50 classifiers (5 feature sets × 10 bootstrap cross-validation iterations). All features are scaled and centered. Finally, we evaluated the clinical meaning of the top transitive sequences used in the phenotyping algorithms.

## RESULTS

As expected, we mined millions of tSPM sequences. Table 1 presents the number of unique representations by type before and after MSMR’s sparsity reduction. On average, we used over 7,000 unique medication and diagnosis codes, from which we mined, on average, over 322,000 SPM and about 6,000,000 tSPM sequential representations. Removing sparse representations (prevalence smaller than 0.5 percent), resulted in on average over 4,000, 28,000, and 1,000,000 unique AVR, SPM, and tSPM features, respectively. Using the mutual information and the joint mutual information filters, the MSMR algorithm further shrunk these features to a final vector of between 50 to 200 features for each phenotype.

To address the research question, phenotyping results are presented in Table 2 (also illustrated in Figure 2). Overall, we found that temporal sequences provided the best phenotyping performances across all 10 phenotypes (except in CAD where we had a tie). Combining sequences with AVR features only resulted in the best overall performance in COPD and RA. Among the sequential representations, in an overwhelming majority of the phenotypes, transitive sequences were included in the best results. The two exception to this were in CAD (in which the difference was 0.001) and T2DM. For six of the 10 phenotypes, we were able to find AUC ROCs reported from other published phenotyping studies. In five of the six phenotypes, the performances we obtained from temporal sequences was substantially superior.

**Table 2.** the area under the receiver operating characteristics curve (AUC ROC)

| data representation | phenotype | AUC<sup>1</sup> | AUC<sup>2</sup> | literature | phenotype | AUC<sup>1</sup> | AUC<sup>2</sup> | literature |
| --- | --- | --- | --- | --- | --- | --- | --- | --- |
| AVR | AD | 0.869 | 0.872 | NA | RA | 0.962 | 0.963 | 0.933-0.961<br>[11,58,59] |
| SPM |  | 0.863 | 0.875 |  |  | **0.970** | 0.970 |  |
| SPM+AVR |  | 0.883 | 0.886 |  |  | **0.970** | 0.971 |  |
| tSPM |  | **0.898\*** | **0.913** |  |  | 0.966 | 0.966 |  |
| tSPM+AVR |  | 0.851 | 0.857 |  |  | 0.964 | **0.972** |  |
| AVR | AFIB | 0.940 | 0.940 | NA | Stroke | 0.875 | 0.876 | NA |
| SPM |  | 0.941 | 0.942 |  |  | 0.880 | 0.881 |  |
| SPM+AVR |  | 0.942 | 0.942 |  |  | 0.887 | **0.894** |  |
| tSPM |  | **0.943** | **0.943** |  |  | **0.879** | 0.879 |  |
| tSPM+AVR |  | 0.940 | 0.941 |  |  | 0.876 | 0.877 |  |
| AVR | CAD | **0.976** | **0.976** | <u>0.896-0.93</u><br>[11,58,59] | T1DM | 0.980 | 0.980 | 0.981<br>[59] |
| SPM |  | 0.974 | 0.974 |  |  | 0.978 | 0.978 |  |
| SPM+AVR |  | **0.976** | **0.976** |  |  | 0.978 | 0.978 |  |
| tSPM |  | 0.975 | 0.975 |  |  | **0.993** | **0.993** |  |
| tSPM+AVR |  | 0.975 | 0.975 |  |  | 0.978 | 0.978 |  |
| AVR | CHF | 0.883 | 0.885 | 0.72-0.87<br>[9,60,61] | T2DM | 0.936 | 0.936 | 0.9<br>[9,59] |
| SPM |  | 0.868 | 0.868 |  |  | 0.944 | 0.945 |  |
| SPM+AVR |  | 0.866 | 0.872 |  |  | **0.956** | **0.959** |  |
| tSPM |  | **0.898** | **0.901** |  |  | 0.926 | 0.926 |  |
| tSPM+AVR |  | 0.893 | 0.896 |  |  | 0.922 | 0.922 |  |
| AVR | COPD | 0.862 | 0.862 | NA | UC | 0.955 | 0.955 | 0.87-0.975<br>[9,30,58,59] |
| SPM |  | 0.866 | 0.867 |  |  | 0.953 | 0.953 |  |
| SPM+AVR |  | 0.864 | 0.866 |  |  | 0.951 | 0.953 |  |
| tSPM |  | 0.860 | 0.860 |  |  | **0.957** | **0.957** |  |
| tSPM+AVR |  | **0.871** | **0.872** |  |  | 0.953 | 0.953 |  |

<sup>1</sup> the median AUC ROC. <sup>2</sup> the top AUC ROC.
\* Top within-phenotype performances are in Bold.

**Figure 2.**
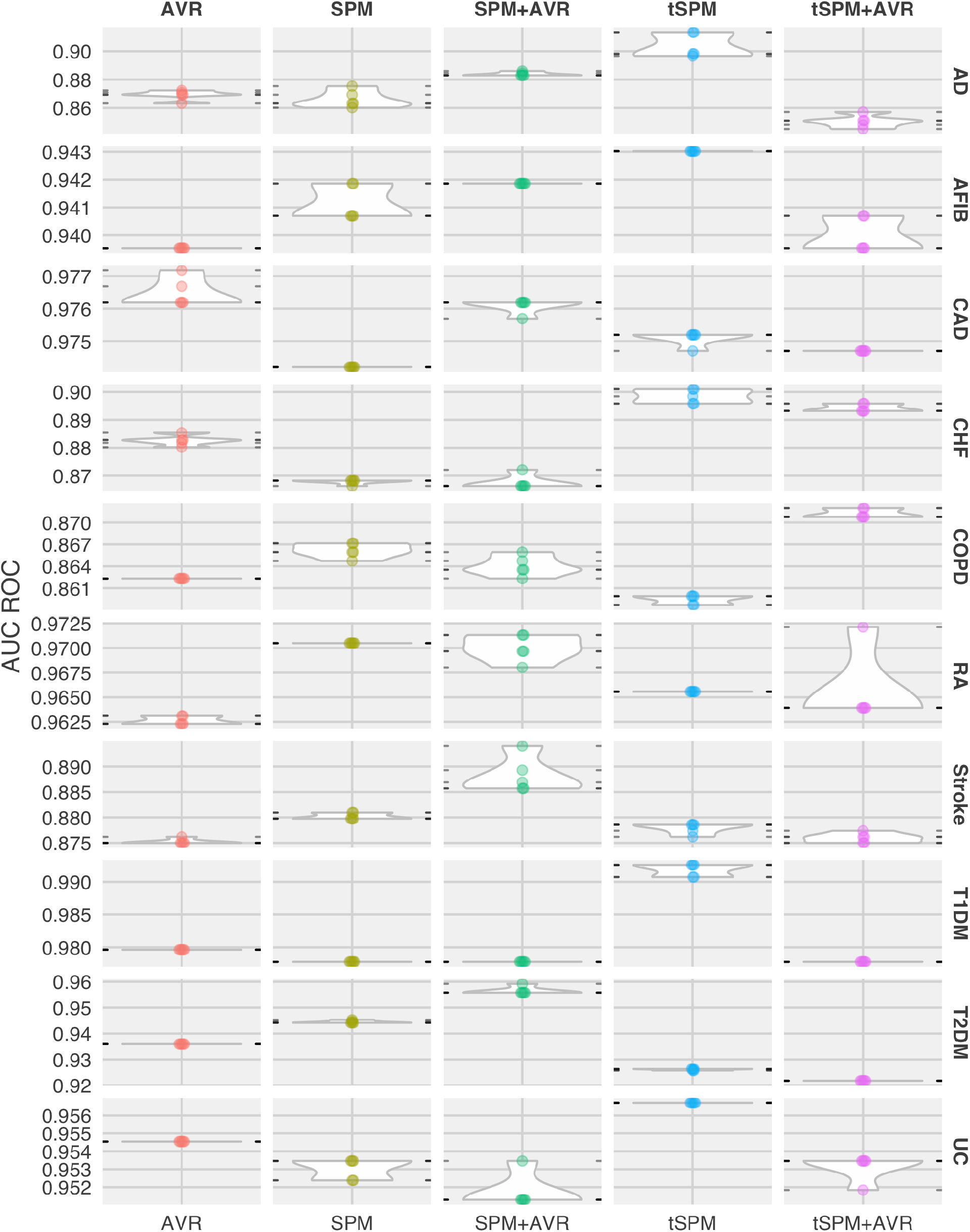
Distribution of phenotyping AUC ROCs by phenotype and data representation.

### Clinical evaluation of the top transitive sequences

From a clinical standpoint, the important transitive sequences can be subdivided into specific categories. Many of the most common sequences were the diagnosis code paired with a risk factor, symptom, complication, or treatment for that disease. It was also common to see the disease code paired with the influenza or pneumococcal 23 vaccination (PPSV23). Table 3 shows a sampling of the identified components that were found to be in sequence with the respective disease for the given phenotype. Each of these components, when in sequence with the diagnosis code, has increased accuracy for identifying the phenotype. For example, in the case of coronary artery disease (CAD), the diagnosis code on its own only identifies true coronary artery disease 34% of the time – i.e., based on the chart reviews, only 34 percent of the patients that had the diagnosis code actually had a confirmed case of CAD. However, when the risk factor, hypertension, precedes the diagnosis code, the accuracy increases to 68%. If the symptom chest pain precedes coronary artery disease the sequence accuracy increases to 70%. If the complication cardiac dysrhythmia, precedes the diagnosis, the accuracy of the sequence increases to 71%. And if the treatment clopidogrel, precedes the diagnosis, the accuracy of the sequence is 97%.There are also cases where the diagnosis code is in sequence with a vaccination or a need for a vaccination. This also leads to increased accuracy for the sequence compared to the components. For example, the sequence “PPSV23 → Rheumatoid arthritis” is 75% accurate for rheumatoid arthritis. Whereas “PPSV23” is only 22% accurate and “Rheumatoid Arthritis” is only 68% accurate.

**Table 3.**
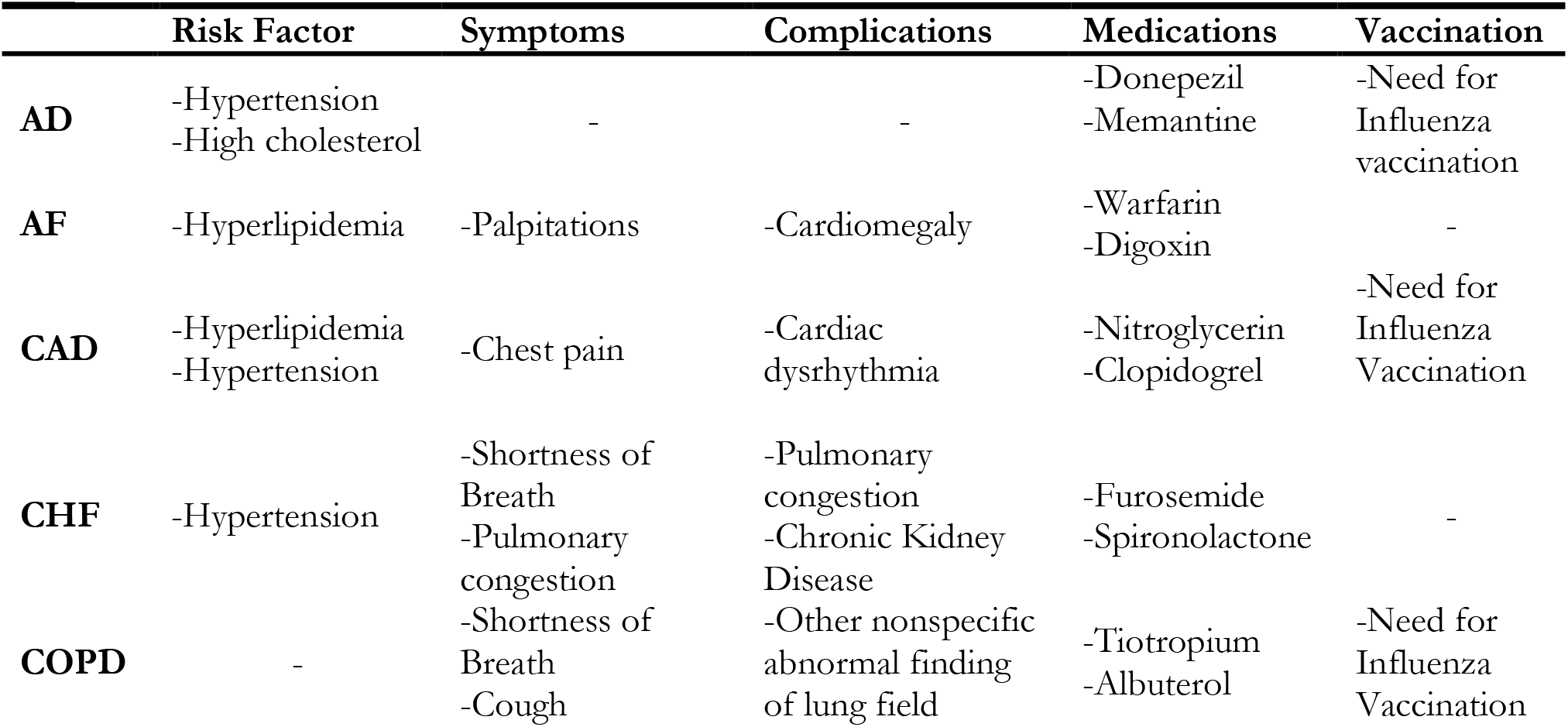
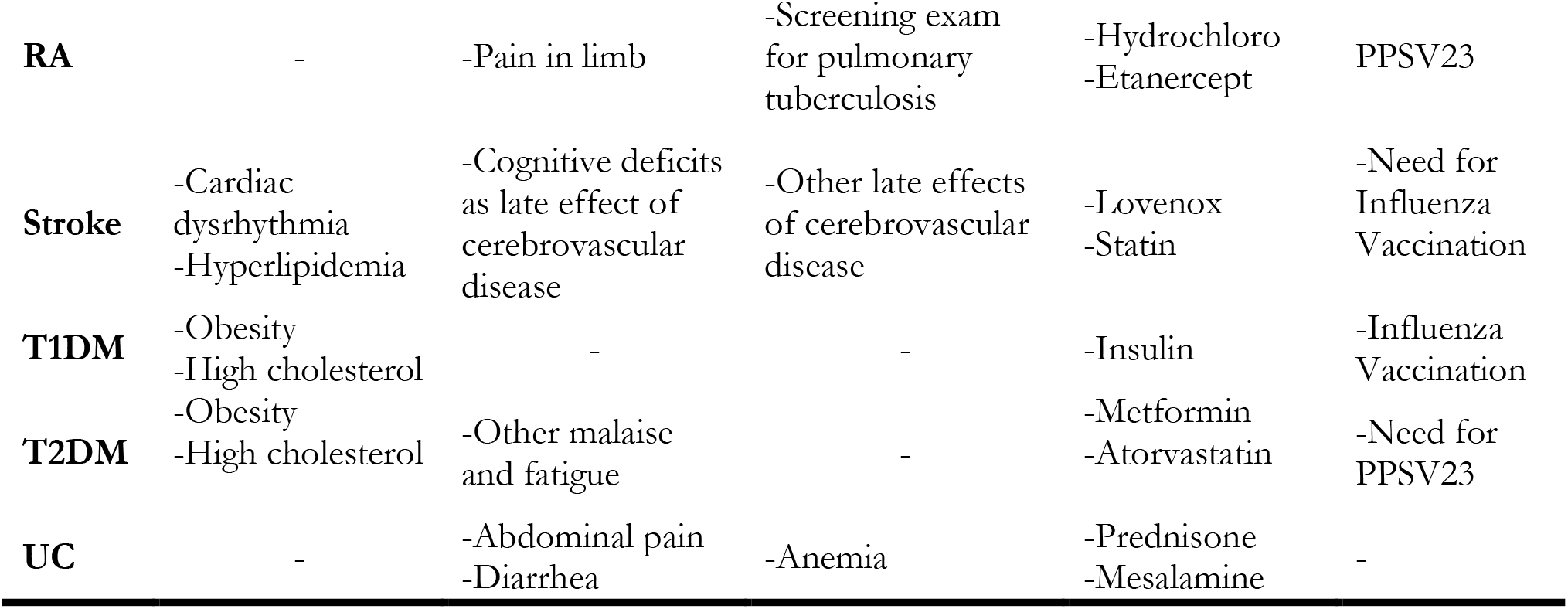
Common components of sequences associated with each phenotype

In some cases, the diagnosis code it not even included in the sequence. Instead the sequence is composed exclusively of risk factors, symptoms, complications, treatments or vaccines, and still offers a high level of accuracy for identifying the specific phenotype. For example, in the case of COPD, “Cough → Tiotropium” is 49% accurate for identifying COPD and only includes a symptom and a treatment. Sometimes both components of the sequence come from the same category. For example, in the case of Diabetes Mellitus Type I, “Insulin → Glucagon” is a sequence of two treatments and accurately identifies the phenotype 63% of the time.

## DISCUSSION

We argue that sequences present more precise information by reducing some of the noise in the EHR data. For instance, we might have *N* observations of the diagnosis code *B* in patient *i*’s medical record. When the diagnosis code *B* is deemed a relevant feature for phenotype *X* (whether through our proposed MSMR or expert ascertion), in the conventional (AVR) approach for computational phenotyping *B* is directly incorporated (as a feature) into the classification algorithm. Sequential data representations, instead, provide a more precise way for incorporating the record *B* into downstream modeling, in that only a proportion of the record *B* may hold useful information for classification that precedes another record (e.g., *B → C*) or follows another record (*A → B*). The MSMR algorithm allows for seamless discovery of such precise sequential record combinations.

Furthermore, the sequences not only lead to more precise phenotyping but they also uniquely capture two distinct events that reflect the patient’s experiences of those events.. For example, “Cardiac dysrhythmia -> Cognitive deficit as a late effect of cerebrovascular disease” is an important sequence for identifying the stroke phenotype. This is likely a common narrative for a stroke patient. The individual develops a cardiac dysrhythmia that leads to the formation of a clot in the heart, that then causes a cerebrovascular accident which leads to cognitive deficits. By sequencing the diagnoses and medications, we developed a rich feature set where individual labels can accurately tell a patient’s story. Analyzing such sequences could give new insight into disease trajectories. Applying this method to new and emerging diseases (such as Covid-19) and then analyzing the sequence features could help us to understand how the disease progresses.

The diagnostic labels and medications listed in Table 3 were all selected by the proposed MSMR algorithm as the significant features for identifying the given phenotype. Each of these labels when in sequence with the disease also have clinical meaning. For example, “Hypertension -> Alzheimer’s Disease”, suggests hypertension precedes Alzheimer’s Disease. The scientific literature supports this relationship. Several longitudinal studies have shown midlife hypertension is consistently associated with the development of Alzheimer’s disease implying that hypertension is a risk factor.[62–64] Another example is the sequence “Congestive heart failure -> Chronic kidney disease”. Again, the literature supports this relationship. In a systematic review of 16 studies with more than 80,000 patients with congestive heart failure, 29% had moderate to severe kidney impairment.[65] Therefore, congestive heart failure preceding chronic kidney disease makes sense because chronic kidney disease is a known complication. These sequences are not just more accurate, but they tell a clinical narrative that corresponds with the patient’s experience.

While risk factors, symptoms, complications and medications paired with the disease may make intuitive sense, vaccinations in sequence with the disease are less obvious. Their inclusion in such sequences may be a result of the specific criteria for receiving the vaccination increasing the probability that the disease code matches the phenotype. The CDC recommends PPSV23 for patients under 65 with chronic diseases such as heart disease, lung disease, and diabetes mellitus and all adults over 65.[66] And while influenza is recommended yearly to all adults, it is very important for those at high risk for serious complications to influenza.[67] The sequence of a vaccination with the disease code may be an accurate label because the vaccination’s presence in the chart further verifies that the patient has a chronic disease.

More complex algorithms such as Recurrent Neural Networks (RNNs)[68] and RNN-based models such as Long Short-Term Memory (LSTM)[69] and Gated Recurrent Unit (GRU)[70] have been used to account for time.[71–78] These algorithms often result in highly predictive models, but they are hard to understand, limiting their utility in healthcare settings. The transitive sequences are similar to simple forms of recurrent events in RNN-based models. The difference is that we do not provide any gate or memory constraint and would accept all possible sequences. This resulted in a large dimensional space. The MSMR algorithm is allowed to pick up what is relevant to the outcome of interest. However, in this paper we only studied 2-deep sequences. We envision extracting deeper sequences, which would further increase dimensionality. In that case, future research may need to apply memory constraints on what to remember from the past.

Despite the billions of dollars that have been spent to institute meaningful use of EHR systems over the past several decades, challenges still remain for using EHR data to rapidly address pressing health issues including the COVID-19 pandemic. This Machine Learning (ML) pipeline, which includes both representation mining and the MSMR algorithm, is capable of engineering predictive features without the need for expert involvement to model different phenotypes and outcomes. Without that bottleneck, this method provides a much faster way for extracting meaningful data from the EHR.

## CONCLUSION

We presented a high-throughput approach for computational phenotyping using temporal sequential data representations. Feature engineering in this approach is fully automated using silver-standard labels. We also demonstrated that using transitive sequences of EHR diagnosis and medication records as features for computational phenotyping yields improved phenotyping performance compared to the timeless raw EHR records. Sequential data representations provide a precise mechanism for incorporating raw EHR records into downstream Machine Learning. Together, the temporal sequences and the ML pipeline can be rapidly deployed to develop computational models for identifying and validating novel disease markers and advancing medical knowledge discovery.

## ACKNOWLEDGMENTS

The authors would like to thank Dr. Sebastien Vasey for invaluable contribution to the MSMR algorithm.

